# On the Number of Driver Nodes for Controlling a Boolean Network to Attractors

**DOI:** 10.1101/395442

**Authors:** Wenpin Hou, Peiying Ruan, Wai-Ki Ching, Tatsuya Akutsu

## Abstract

It is known that many driver nodes are required to control complex biological networks. Previous studies imply that *O*(*N*) driver nodes are required in both linear complex network and Boolean network models with *N* nodes if an arbitrary state is specified as the target. In this paper, we mathematically prove under a reasonable assumption that the expected number of driver nodes is only *O*(log_2_ *N* + log_2_ *M*) for controlling Boolean networks if the targets are restricted to attractors, where *M* is the number of attractors. Since it is expected that *M* is not very large in many practical networks, this is a significant improvement. This result is based on discovery of novel relationships between control problems on Boolean networks and the coupon collector’s problem, a well-known concept in combinatorics. We also provide lower bounds of the number of driver nodes as well as simulation results using artificial and realistic network data, which support our theoretical findings.

## I. Introduction

Boolean network (BN) is a sequential dynamical system composing of a large number of highly interconnected processing nodes the states of which are updated by Boolean functions of other nodes and/or itself [1]. It is simple but very efficient in modeling genetic regulation [2–4], neural networks [5], cancer networks [6], quorum sensing circuits [7], cellular signaling pathways [8], dynamic games [9], computer design [10], and social networks [11]. Each node in a BN takes either 0 or 1, where 0 and 1 mean the node is inactive and active, respectively. Since a *BN* with *N* nodes has 2^*N*^ possible states, it will eventually reach a previously visited state, thus stay in that state circle, called an attractor.

Among various problems on BNs, control of a BN is particularly important in which the values of a subset of nodes or external signals are manipulated so as to drive the BN to a desired state [9, 12–16]. For example, in disease treatment, one may need to conduct therapeutic intervention that drives the cell state of a patient from a current state to a desired state such as a benign state, and keep this state afterward. Interventions can be additional drugs, hormones, or the embedded genes that can be manipulated as the control variables of the networks [16]. However, it is difficult to give many drugs or to manipulate many genes in practice. Therefore, it is important to select a small subset of nodes from the nodes in a given BN so that the BN is driven to a desired state by manipulating the values of these nodes only. These nodes are called the *driver nodes*.

Extensive studies have been done on control of BNs [15, 17–20], control of complex networks with linear dynamics [21–23], and the chariterization of multiagent controlability from network structures [24, 25]. Recently, semi-tensor product (STP) proposed in [13, 18] has been used widely in various control models of BN [16, 26–31]; the stability and stabilization have also been considered as an important aspect in control of BNs [32–36]. However, it is known that *O*(*N*) driver nodes are required to control linear complex networks [21] if an arbitrary state is specified as the target, although a single driver node is enough if the network has a special structure (every node has a self loop) [37]. Another approach has recently been proposed for control of BNs by restricting the target states to attractors (see also Fig. 1). Posing such a restriction is reasonable because target states are stable states in many practical cases, for example, healthy and stable states in disease treatment. Mochizuki et al. showed that the Feedback Vertex Set (FVS) can be a set of driver nodes if the targets are attractors [38], where the relationship between singleton attractors and FVS was previously found [39, 40]. Zañudo and Albert [41] proposed a method to identify driver nodes for attractors of a BN irrespective of the updating scheme (synchronous/asynchronous) and the number of time steps with stable motifs each of which is defined as a set of nodes that forms a minimal strongly connected component of the network. Concerning the scaling behavior of the minimum control cost of BNs without structural or temporal constraints, however, there is not yet a conclusive results in literature. While simulation results suggest that a small number of driver nodes are enough if the target states are restricted to attractors [38, 41], results from [14, 20] imply that *O*(*N*) driver nodes are required if an arbitrary state is specified as the target.

**Fig. 1:**
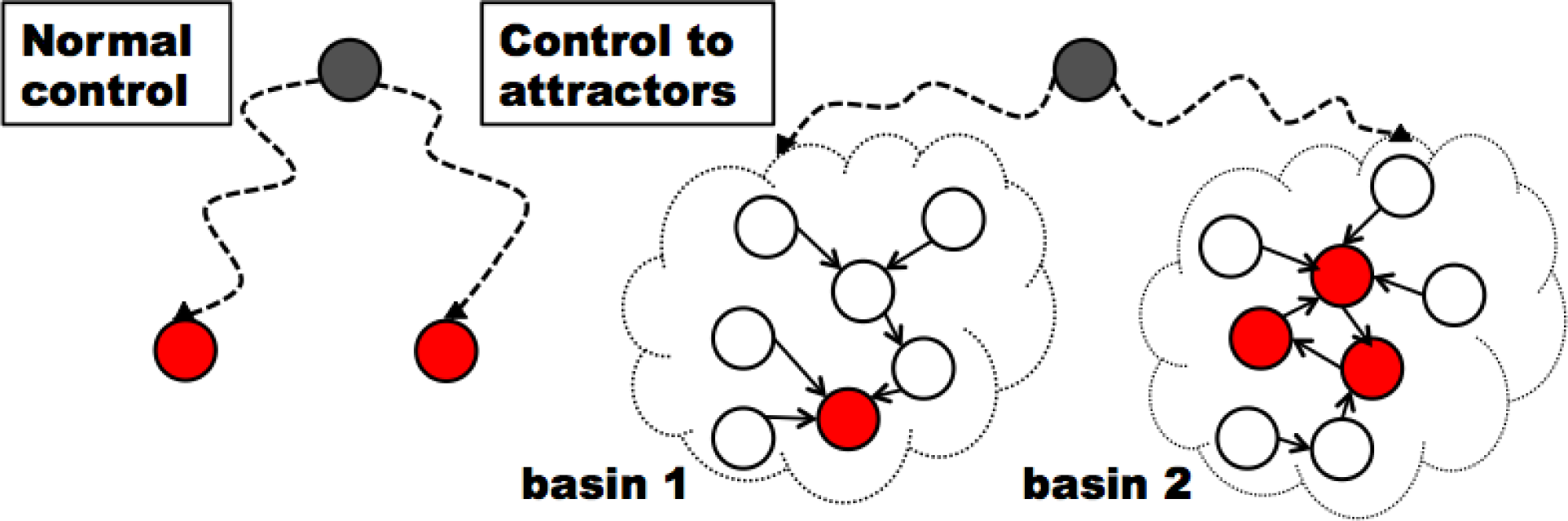
Comparison of Normal Control and Attractor-based Control. Circles denote states where grey ones are initial states and red ones are targets.

In this paper, we mathematically prove that the expected number of driver nodes is only log_2_(*N*) + log_2_(*M*) + 2 if the targets are restricted to attractors, under a reasonable assumption. Since it is expected that *M* is not very large in many practical networks, this is a significant improvement. Even if the number of attractors, *M*, grows as 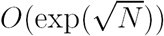 [42], it is still 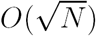, which is much smaller than *O*(*N*). Our theoretical analyses are based on a discovery of novel relationships between BN control problems and Coupon Collector’s Problem, a well-known concept in combinatorics. Based on these relationships, we also prove that the expected number of driver nodes is ln(*M* ln *M*) + ln 3 (resp., log_2_(*M* ln *M*) + 2) if both initial and target states are specified (resp., only a target state is specified). Note that these numbers are very small. For example, the former number is around 8.06 for *M* = 200. Since attractors are often regarded as cell types and the number of cell types in human is often said to be aounnd 200 [43], these results suggests that control of a small number of genes is enough to modify types of cells. The differences on three bounds show a nontrivial nature of mathematical analyses although all results are based on relationships with BN control problems and Coupon Collector’s Problem. Furthermore, we provide lower bounds of the number of driver nodes as well as simulation results supporting our theoretical findings. In particular, we show that the derived upper bounds are close to the lower bounds in the average case, whereas the lower bounds are much higher if we consider the worst case.

## II. Preliminaries

### A. Boolean Networks

A *Boolean network G*(*V, F*) consists of a set of *N nodes V* = {*v*_1_*, …, v*_*N*_ } and a list of *Boolean functions F* = (*f*_1_*, …, f*_*N*_). The state of *v*_*i*_ at time *t* is denoted by *v*_*i*_(*t*). The vector **v**(*t*) = (*v*_1_(*t*)*, …, v*_*N*_ (*t*)) denotes the *state* of the BN at time *t*. The *Boolean function* for node *v*_*i*_ is a logical combination of *k*_*i*_ (called *in-degree*) variables in form of *f*_*i*_(*v*_*i*1_*, …, v*_*ik*_*i*) where *v*_*i*1_, *…*, *v*_*ik*_*i* are *k*_*i*_ input nodes of *v*_*i*_. Then the state of node *v*_*i*_ at time *t* + 1 is *v*_*i*_(*t* + 1) = *f*_*i*_(*v*_*i*1_(*t*)*, …, v*_*ik*_*i* (*t*)), or equivalently, *v*_*i*_(*t* + 1) = *f*_*i*_(**v**(*t*)). The state of the (synchronous) BN is determined by **v**(*t* + 1) = **f** (**v**(*t*)). If **v**(*t* + *r*) = **f** ^*r*^(**v**(*t*)) = **v**(*t*), then {**f** ^*j*^(**v**(*t*))} (*j* = 1*, …, r*) is called an *attractor* of length *r*. Additionally, the *basin* of an attractor is defined as the set of states leading the system to the attractor. We denote the number of basins (i.e., the number of attractors) by *M*, and the set of states in the *i*th basin by *A*_*i*_. Starting from any initial state, a BN eventually falls into one of its attractors (Fig. 2). For two states **v**_*i*_ and **v**_*j*_, *d*(**v**_*i*_, **v**_*j*_) denotes the Hamming distance between **v**_*i*_ and **v**_*j*_ (i.e., the number of distinct bits).

**Fig. 2:**
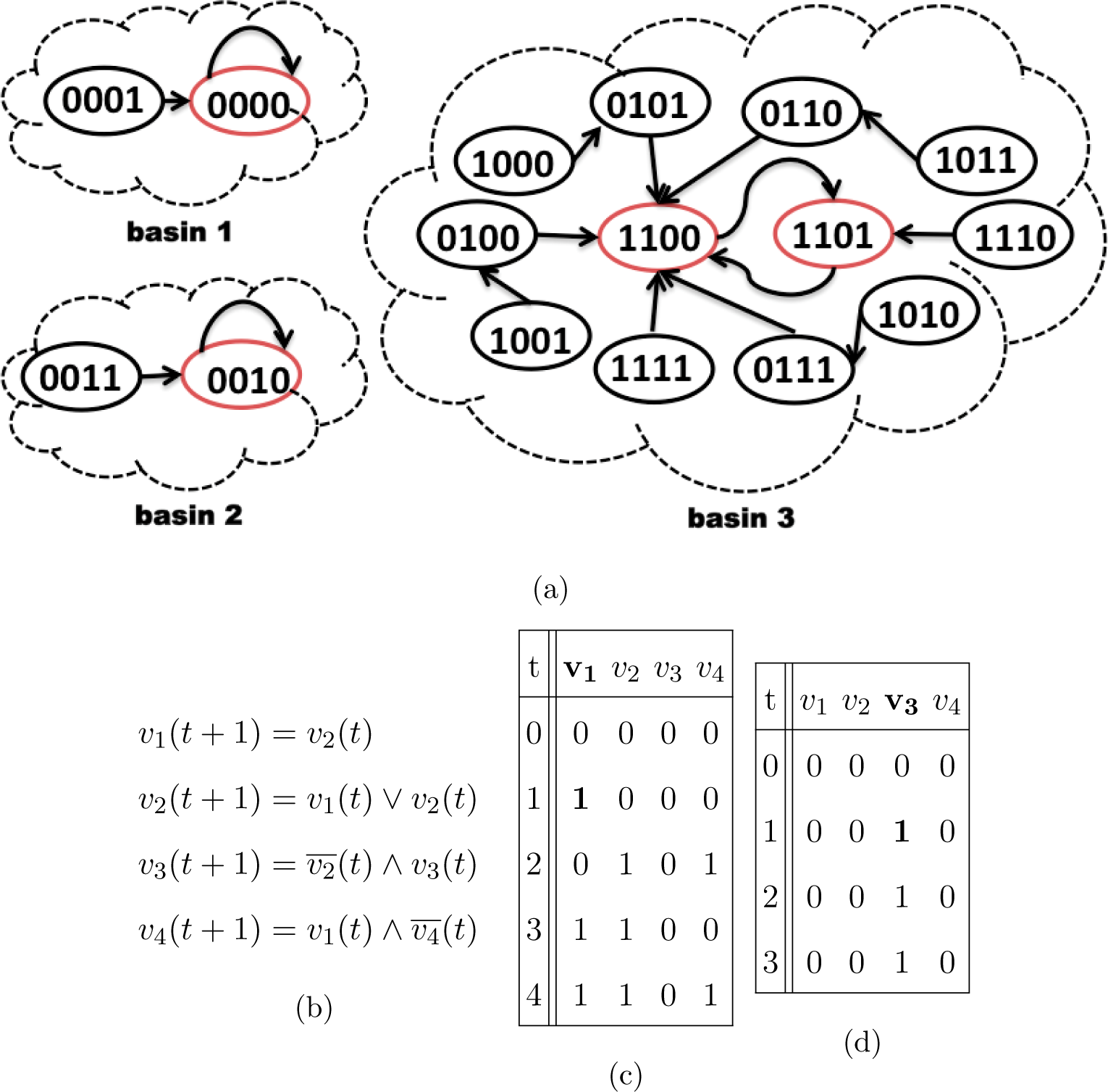
Examples of a BN and its control. (a) Basins & state transitions. (b) Boolean transition rules. (c) BN control: state 0000 to basin 3. (d) BN control: state 0000 to basin

### B. Problem Definitions

We consider the *minimum driver set problem* for a BN [14], which is defined as follows. We are given a BN, an initial state **v**^0^, and a target state **v**^*T*^, where the target time step may or may not be given. Then, a subset *U* of *V* is called a set of *driver nodes* (a driver set, for short) if there exists a sequence of states of *U* that drives the BN from **v**^0^ to **v**^*T*^ (if the time step is specified, the BN must take state **v**^*T*^ at the specified time step). If the number of elements in *U* is the minimum among such sets, *U* is called the *minimum driver set*. The task is to find this minimum driver set. Note that there always exists a solution: if we let *U* = *V*, *U* is a driver set (although it is not necessarily the minimum). A sequence of states given to *U* corresponds to a sequence of control signals.

In this paper, we focus on the case where *the target state is any state in the specified attractor and the control is given only at time step 1*. Then, it is enough to drive a BN to any state in the target basin since the BN then falls into the target attractor. For example, in Fig. 2(c), the initial state is 0000 and the target state is 1101. If we let *v*_1_(1) = 1, the BN reaches 1101 at *t* = 4 and stays in the third attractor. In Fig. 2(d), the initial state is 0000 and the target state is 0010. If we let *v*_3_(1) = 1, the BN reaches 0010 at *t* = 1 and stays in the second attractor. In both cases, we need to change the state of 1 node (i.e., 1 bit), where {*v*_1_} and {*v*_3_} correspond to the minimum sets of driver nodes, respectively. Therefore, the number of driver nodes is 1 in both cases. However, if we are requested to drive the BN to any attractor using the same set of driver nodes, the minimum set of driver nodes is {*v*_1_*, v*_3_}. In these examples, we assumed that the state at *t* = 1 is determined only by flipping some bits in a driver node set, for the sake of simplicity. However, all the results in this paper hold if we modify the formulation so that state transition and control are applied at the same time step, where application of a control is limited to step 1.

As mentioned above, a minimum driver set may or may not depend on a target state. Similarly, it may or may not depend on an initial state. Therefore, we consider the following three problems.

**Problem 1** [Attractor-dependent Control]

Instance: a BN with *N* nodes, an initial state, a target basin.

Output: a minimum set of driver nodes with which we can steer the BN to the basin in one control.

**Problem 2** [Attractor-independent Control]

Instance: a BN with *N* nodes, an initial state.

Output: a minimum set of driver nodes with which we can steer the BN to any basin of the BN in one control.

**Problem 3** [Anycast Control]

Instance: a BN with *N* nodes.

Output: a minimum set of driver nodes with which we can steer the BN from any initial state to any basin of the BN in one control.

### C. Coupon Collector’s Problem

*Coupon Collector’s Problem* (CCP) is a random allocation problem in probability theory describing "collect all coupons and win" game. Assume there are *M* coupons in an urn, each with a probability *P*_*i*_(*i* = 1*, …, M*) to be collected. In each trail, you can draw one coupon, and then put it back to the urn. CCP describes at least how many trials are needed to have *i* collections (*i* different coupons have been collected in history). If *i* = *M*, then you achieve a full collection.

## III. Theoretical Results

### A. Upper Bounds via Coupon Collector’s Problem

We analyze three problems independently and then compare their results. Our common idea is that the structure of a BN has already been embedded in the generation of the basins of attractors, therefore by considering driving the BN to basins, we take into account the topology and dynamic processes together tactfully. Our theoretical results are based on a discovery of novel relationships between the three problems and CCP, each in a different way. Basically, we associate each state of a BN to a type of coupon so that states in the same basin have the same type, and then analyze how many bits should be flipped in order to have a specified coupon or all coupons. However, it is difficult to analyze three cases in a unified way and thus we consider three cases separately.

Although we assume in most parts of theoretical analysis that the basins are obtained by a random partition of 2^*N*^ states into *M* sets of the same size, the restriction of the size will be considerably relaxed, to be stated in Corollary 1.

#### Theorem 1

*Assume that the basins are obtained by a random partition of* 2^*N*^ *states into M sets of the same size. Then, for M* ∈ [3, 1.44^*N*^] *with sufficiently large N, the expected minimum number of driver nodes in Attractor-dependent Control is bounded above by* ln(*M* ln *M*) + ln 3.

##### Proof

We formulate the Attractor-dependent Control problem as CCP. Let **v**^0^ be the initial state. Without loss of generality, we assume **v**^0^ is a vector of bits whose values are all 0. Let 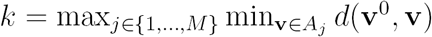. It means that there exists at least one state with at most *k* bits value 1 in each basin and thus this *k* gives the minimum number of driver nodes in Attractor-dependent Control.

We sort the *N* -bit states in the following order (i.e., increasing order of the number of 1s):

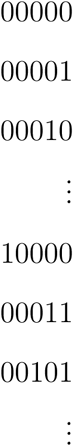

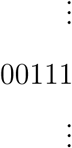

Then, each vector is considered as a coupon belonging to one of *M* types where the type is essentially differentiating which basin that state belongs to. We draw coupons in the above order. Suppose that after drawing the *h*th coupon, we have all *M* types. Then, the number of 1s in the *h*th coupon gives *k*.

It is well-known that the expected number of trials to have a full collection of all *M* types of coupons is

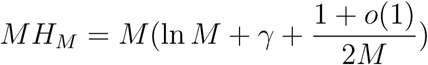

if all types are being collected equally likely [44]. Here *H*_*M*_ is the *M* th harmonic number, *γ* ≈ 0.5772156649 is the Euler-Mascheroni constant. When *M* ≥ 2, we have

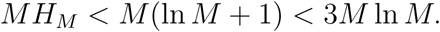

Note that the probability of drawing a coupon of a new type is increasing actually because we draw coupons by following the above list without replacement. As a result, the number of trials needed to have a full collection is actually smaller than *MH*_*M*_, which further indicates that we are deriving an upper bound.

Here we recall that there exist *M* basins. Although one basin is specified in Attractor-dependent Control, one of the elements of this basin must be included in the set of states with at most *k* 1s if the size of a driver set is *k* (since we are assuming that **v**^0^ is the zero vector). Furthermore, in order to have at least one state from any specified basin, 3*M* ln *M* states are needed in the average case. Since the number of states with at most *k* 1s is

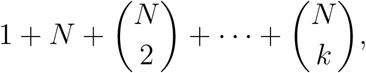

we have a required driver set if

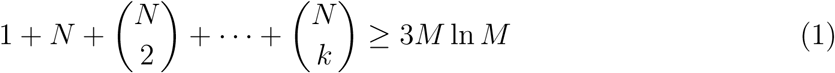

holds. Of course, the expected number of driver nodes does not exactly correspond to the expected number of coupons. However, since the probability that the number of trials needed to cover all coupons is larger than 3*M* ln *M* is quite small [45], it is enough that InEq. (1) is satisfied.

Since 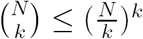 holds, InEq. (1) is satisfied if the following is satisfied

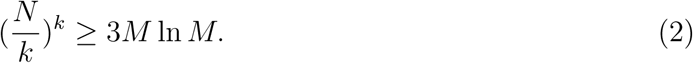

By taking logarithm of both sides of this inequality, we have

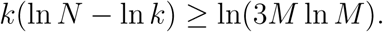

Here, we assume 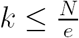, where *e* is the base of the natural logarithm. Then, we have

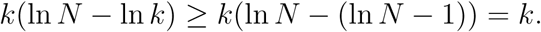

Therefore, InEq. (2) (and thus InEq. (1)) is satisfied if

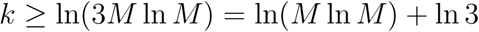

is satisfied.

In order to ensure the existence of such *k*,

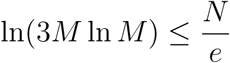

must be satisfied because we assumed 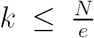. Suppose that *M* < *α*^*N*^ holds for some *α* < *e*^(^1^/*e*)^. Then, we have

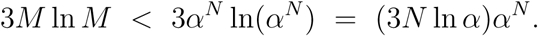

Since (3*N* ln *α*)*α*^*N*^ < (*e*^(^1^/*e*)^)^*N*^ holds for sufficiently large *N* for any positive constant *α* < *e*(1/*e*),

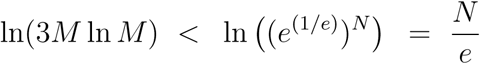

holds for sufficiently large *N* for any positive constant *α* < *e*^(^1^/*e*)^. Therefore, the theorem follows from 1.44 < *e*^(^1^/*e*)^.

#### Theorem 2

*Assume that the basins are obtained by a random partition of* 2^*N*^ *states into M sets of the same size. Then the expected minimum number of driver nodes in Attractor-independent Control is bounded above by* log_2_(*M* ln *M*) + 2.

##### Proof

As in the proof of Theorem 1, we formulate the Attractor-independent Control problem as CCP. Since we need to use a fixed set of driver nodes for all attractors, we consider the different ordering of the *N* -bit states. We sort these states in the increasing order of their values (see also Fig. 3):

**Fig. 3:**
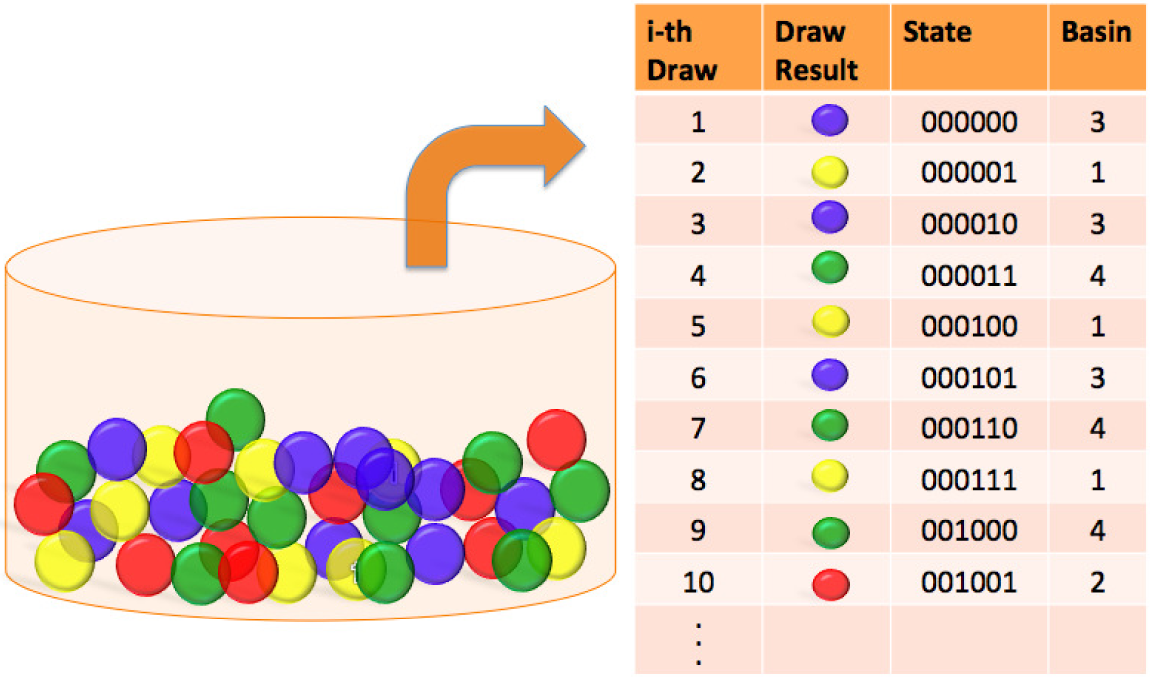
Attractor-independent Control formulated as CCP: *M* = 4 as an example.

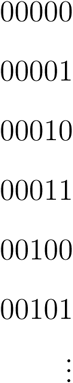

Again, each vector is considered as a coupon belonging to one of *M* types. We draw coupons in the above order. Let *τ* be the number of states needed to collect all *M* labels. We can see from the ordering that the first *N -*⌈log_2_(*τ*) ⌉ bits of all vectors until *τ* are all 0, which means that it is enough to control the last ⌈log_2_(*τ*) ⌉ bits. As in the proof of Theorem 1, the expected number of *τ* is upper bounded by 2*M* ln *M*. Since 𝔼[log_2_(*X*)] ≤ log_2_(𝔼[*X*]) holds from Jensen’s inequality, we have

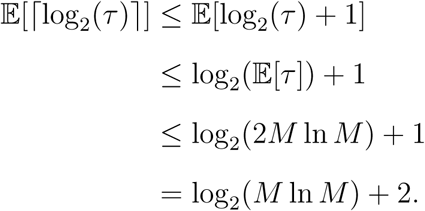

#### Theorem 3

*Assume that the basins are obtained by a random partition of* 2^*N*^ *states into M sets of the same size. Then, for M* < 1.587^*N*^ *, the expected minimum number of driver nodes in Anycast Control is bounded above by* ⌈log_2_(*N*) + log_2_(*M*) + 2 ⌉ *for sufficiently large N*.

Suppose all 2^*N*^ states are balls in an urn (left). Colors yellow, red, blue, green of the balls are labels showing the ball belongs to basin 1,2,3,4, respectively. In each trial, we draw a ball, record its label, and then put it back to the urn. Draw results are shown in the table (right). It shows that in the tenth draw, it is the first time we have drawn balls from all 4 basins in history. Given **v**^0^, we can drive **v**^0^ to any of these 10 states in one control by the first ⌈log_2_(10)*l* bits of **v**^0^. Notice that ⌈log_2_(10) ⌉ < log_2_(*M* ln *M*) + 2 with *M* = 4 which is consistent with Theorem 2.

##### Proof

We divide *N* bits to the first *H* bits and the remaining *L* bits, where the last *L* bits are used for control (*H* + *L* = *N*). For any bit vector (i.e., state of a BN) **v** of size *N*, **v**_*H*_ and **v**_*L*_ denote bit vectors consisting of the first *H* bits and the last *L* bits, respectively. Namely, **v** = **v**_*H*_ **v**_*L*_. Although this partition is similar to that of the proof of Theorem 2, we cannot assume that the first *H* bits are all 0 since we are considering arbitrary initial states.

If every basin *A*_*i*_ contains states whose first *H* bits cover all 2^*H*^ bit patterns (i.e., (∀*A*_*i*_)(|{**v**_*H*_|**v** ∈ *A*_*i*_}| = 2^*H*^)), we can drive any initial state to any basin. Therefore, we determine *H* so that this property (denoted by property (#)) holds with high probability.

Let *T* be the number of trials needed to collect all *M* coupons. It is known that *T* > *βM* ln *M* holds with probability at most *M* ^*−β*+1^, if each type of coupon is selected uniformly at random (see Section 3.6 of [44]).

We relate Anycast Control with CCP. Different from the previous problems. we regard the first *H* bits of each state as a type of coupon. If each basin contains all 2^*H*^ type coupons, (#) holds.

Suppose that |*A*_*i*_| ≥ 3 *·* 2^*H*^ ln(2^*H*^) holds, where *H* and *L* are determined so that this condition is satisfied. We regard the first 3 *·* 2^*H*^ ln(2^*H*^) vectors of each *A*_*i*_ as coupons. Then, the probability that there is a missed coupon in a basin *A*_*i*_ is at most (2^*H*^)^*−β*+1^ = 2^-2*H*^, where *β* = 3. Since there exist *M* basins, the probability that (#) does not hold is at most

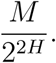

Hereafter, we assume *M* < 2^*−αN*^ and *H* > (1 - *β*)*N* (i.e., *L* < *βN*), where *α, β* > 0 are to be determined later. Then, we have

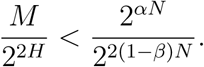

If (#) is not satisfied, we use all *N* bits as driver nodes. However, contribution of such a factor to the expected number is less than

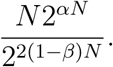

Therefore, we focus on the case where (#) holds.

In order to satisfy the assumption of |*A*_*i*_| ≥ 3 *·* 2^*H*^ ln(2^*H*^), the following inequality must be satisfied:

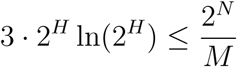

since |*A*_*i*_| = 2^*N*^ /*M* holds. By taking logarithm of both sides, we have

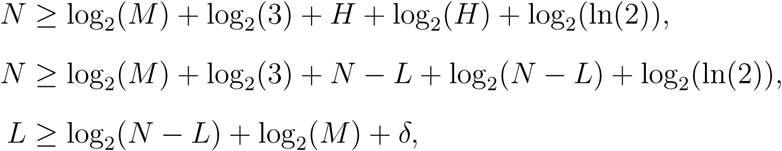

where *δ* = log_2_(3) + log_2_(ln(2)) < 2. The last inequality holds if the following holds:

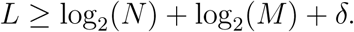

Therefore, the number of driver nodes is bounded above by

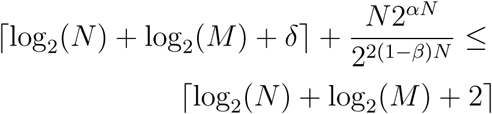

for sufficiently large *N* if the following are satisfied

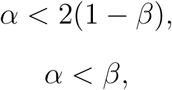

where *α* < *β* comes from ⌈log_2_(*N*) + log_2_(*M*) + 2⌉ < *βN*. By solving 2(1 - *β*) = *β*, we have *β* = 2/3 and *α* < 2/3. Since 2^2^/^3^ > 1.587, we have the theorem.

Here we note that ln(*M* ln *M*) + ln 3 < log_2_(*M* ln *M*) + 2 < ⌈log_2_(*N*) + log_2_(*M*) + 2⌉ holds for *M* ≤ 2^*N*^. This is consistent with the difficulties of the three problems.

Although we assumed in the above that all basins have the same size, this restriction can be considerably relaxed by adding a constant factor. We show this result only for Anycast Control since it is most general.

#### Corollary 1

*Assume that the basins are obtained by a random partition of* 2^*N*^ *states into M sets where each basin has size at least* 2^*N*^ /(*cM*) (*c* ≥ 1)*. Then, for M* < 1.587^*N*^ *, the expected minimum number of driver nodes in Anycast Control is bounded above by* ⌈log_2_(*N*)+ log_2_(*M*) + 2 + log_2_(*c*) ⌉ *for sufficiently large N*.

##### Proof

In the proof of Theorem 3, we assumed that the size of each basin is 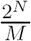 and thus we had a condition of

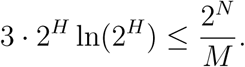

Here, we replace 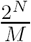 with 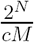 where *c* ≥ 1. Then, the following condition must be satisfied:

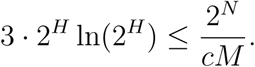

As in the proof of Theorem 3, we have

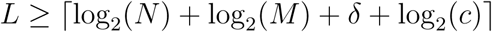

from which the corollary follows.

### B. Lower Bounds

We can also show lower bounds of the number of driver nodes for all three problems. For Attractor-independent Control and Anycast Control, log_2_(*M*) is a trivial lower bound because at least log_2_(*M*) bits are required to differentiate *M* basins.

#### Proposition 1

*For any BN with M attractors, the number of driver nodes in Attractor-independent Control and Anycast Control is bounded below by* log_2_(*M*).

For Attractor-dependent Control, we can change driver sets depending on target attractors. Therefore, we need another analysis method.

#### Proposition 2

*For any BN with M attractors, the number of driver nodes in Attractor-dependent Control is bounded below by* log_*N*_ (*M*).

##### Proof

Let **v**^0^ be the initial state. Without loss of generality, we assume **v**^0^ is a vector of bits whose values are all 0. Let 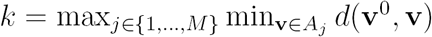. It means that there exists at least one state with at most *k* bits value 1 in each basin. Then the number of such states is bounded above by

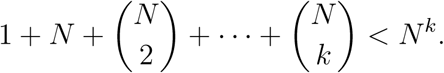

Since the number of states can be produced by driver nodes must be larger than the number of basins to make it possible to drive **v**^0^ to all basins, we have

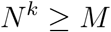

or equivalently

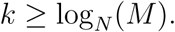

These results suggest that the bounds shown in Theorems 1, 2, and 3 cannot be significantly improved. Recall that it is assumed in all three problems that control is given only at *t* = 1.

In the above, we considered the expected number of driver nodes. If we consider the worst case, a much larger number of driver nodes are required as shown below.

#### Proposition 3

*Suppose that N is even. Then, the minimum number of driver nodes in Attractor-dependent Control is lower bounded by* (*N*/2) + 1 *in the worst case even if M* = 2 *and two basins are of equal size*.

##### Proof

We partition the set of 2^*N*^ states into two basins *A*_1_ and *A*_2_ such that *A*_1_ consists of states each of which has at most *N*/2 1s and consequently *A*_2_ consists of states each of which has at least *N*/2 + 1 1s. Suppose that the initial state consists of *N* 0s (i.e., every bit is 0) and the target basin is *A*_2_. Then, the minimum Hamming distance between the initial state and the states in *A*_2_ is (*N*/2) + 1. Therefore, the proposition holds.

Since Attractor-dependent Control is the easiest among the three problems, this worst case lower bound holds also for Attractor-independent Control and Anycast Control. It is to be noted that the number of driver nodes is trivially upper bounded by *N*. Therefore, this proposition justifies the assumption of considering the average case.

### C. Max-min Analysis Method

We can analyze the minimum number of nodes in Attractor-independent Control without using CCP and can get a more accurate bound if we adopt an assumption that each basin is an *independently selected multi-set* of size 2^*N*^ /*M* whose elements are randomly selected from {0, 1}^*N*^. It follows from this assumption that 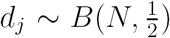, ∀*j* = 1*, …, M*, where *d*_*j*_ is the Hamming distance between a randomly selected initial state **v**^0^ to a state in the *j*th basin, and *B*(*N, p*) denotes the binomial distribution for *N* trials with success probability *p*. To be shown later, this assumption is supported by computer experiments, and the resulting bound better explains the results of computational experiments.

#### Theorem 4

*The expected minimum number of driver nodes of Attractor-dependent Control is* 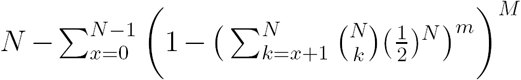 *, if each basin is an independently selected multiset of size* 2^*N*^ /*M whose elements are randomly selected from* {0, 1}^*N*^.

##### Proof

Recall that 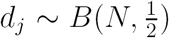. By assumption, 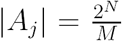, denoted as *m*. Then *d*(**v**^0^, **v**) ∀**v** ∈ *A*_*j*_ is *m* times repetitive binomial trials. Let the results of all trials be 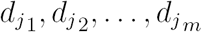 and the order statistics be

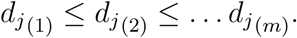

The first order statistics 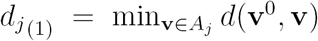. Its cumulative distribution function (CDF) is given by

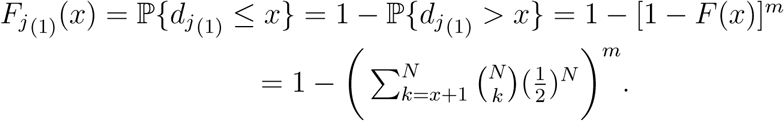

Therefore, for *d*_*j*_ where *j* = 1*, …, M*, we have *d*_1(1)_*, …, d*_*M* (1)_. For convenience, we denote these random variables as *ξ*_1_*, …, ξ*_*M*_. Their order statistics is given by

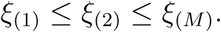

Then the last order statistic *ξ*_(*M*)_ = max_*j*_ min_**v***∈A*__*j*_ *d*(**v**^0^, **v**). Its CDF is given by

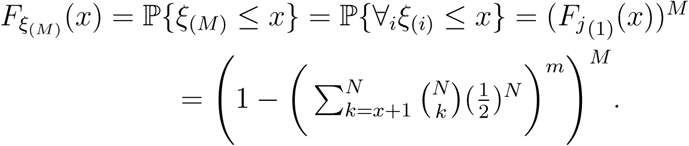

Note that we can calculate the expected value of a non-negative random variable *X* by using its CDF *F* (*X*),i.e., 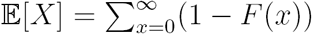 because

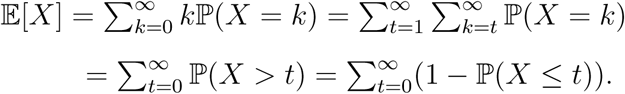

Therefore, we can further give the expected number of 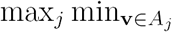 *d*(**u**, **v**) as

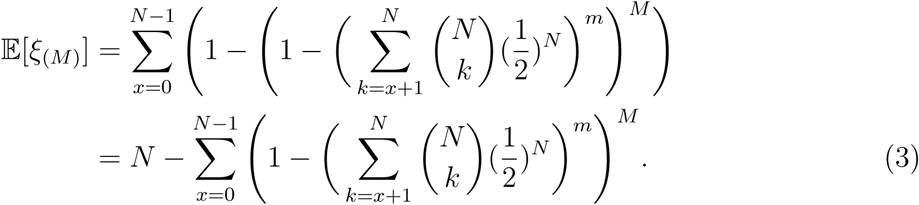

### D. Non-Uniform Basin Size Distributions

We can extend our result on Attractor-independent Control for arbitrary basin size distributions by utilizing a known result on CCP with arbitrary probability distributions [46]. It has been proved in [46] that the expected number of trials *C*_*M*_ needed to have a full collection of *M* different coupons is

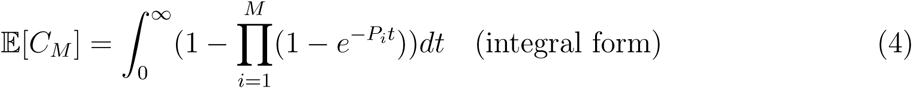

or

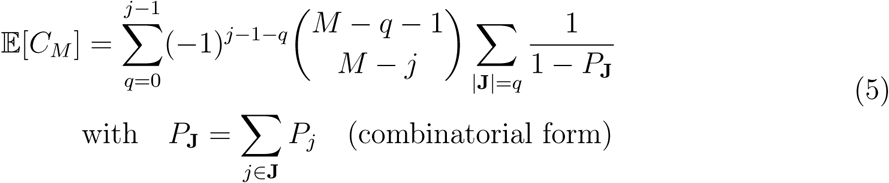

where *P*_*i*_ is the probability of collecting the *i*th coupon, **J** *∈* Ω, and Ω = {1, 2*, …, M* }. By using this result, we have:

#### Theorem 5

*The expected minimum number of driver nodes in Attractor-independent Control for an arbitrary basin distribution is bounded above by* log_2_(𝔼[*C*_*M*_]) + 1.

##### Proof

We associate the minimum of driver nodes in Attractor-independent Control to CCP as in the proof of Theorem 2, with letting *P*_*i*_ = |*A*_*i*_|/2^*N*^. Then, in the same way, the expected minimum number of driver nodes needed to drive the BN to any one basin is bounded above by log_2_(𝔼[*C*_*M*_]) + 1.

Since “+1” factor in log_2_(𝔼[*C*_*M*_]) + 1 is rather an overestimation, we use log_2_(𝔼[*C*_*M*_]) in computational experiments.

To be discussed in Section IV, we empirically find that the distribution of the basin size follows a power-law. Therefore, we consider the case where the basin size follows a power-law.

It has been shown in Example 4.4 in [47] that if the probability of picking the *i*th coupon follows a power-law *i*^*−b*^ (*b* > 0), the expected number of trials needed for a full collection is

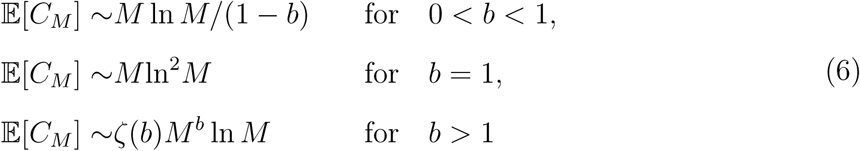

by letting *r* in [47] equal to 1, where *ζ*(*·*) denotes the Riemann zeta function.

It has been shown that any process generating Zipf rank distribution would also have a power-law probability density function (see Appendix 2 in [48]) as outlined below. The relationship between these two is that if **X** follows a power-law ℙ[**X** = *x*] ∼ *x*^*−a*^ then it would also have the *r*th ranked variable *X*_*r*_ satisfying 𝔼[*X*_*r*_] ∼ *Cr*^*−b*^ where *a* = 1+(1/*b*). Therefore, we can apply Eq. (6) to the case when the basin size follows power-law distribution ℙ[**X** = *x*] = *Cx*^*−a*^.

Then, we can apply the analysis method used in the proof of Theorem 2 and obtain the following upper bounds:

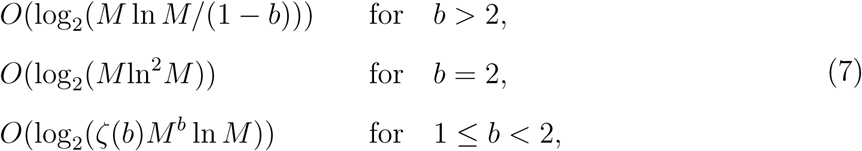

where *b* = 1/(*a -* 1).

## IV. Computational Experiments

### A. Data

The BNs we used for simulation comprise of random BNs and realistic BNs. Random BNs are generated by using C and R programming languages, including random NK-BNs (BNs with *N* nodes and the in-degree is bounded by *K*) and random scale-free BNs with *γ* in different values. There are four realistic BNs constructed from real-valued gene measurements: *yeastTS net* is a BN with two attractors constructed from real-valued time series data of four preselected genes from the yeast cell cycle [49]; *cellcycle net* is a BN with 10 genes and two attractors constructed from mammalian cell cycle network introduced by [3]; *budding net* is a BN model of the control of the budding yeast cell cycle regulation from [50] with 12 genes and seven attractors; *flower net* is a BN model of the control of flower morphogenesis in Arabidobis thaliana from [51] with 15 genes and 10 attractors.

### B. State and Basin Size Distributions

In Theorem 4, we assumed that the distance between an initial state and the states in each basin follows a binomial distribution: 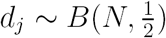, ∀*j* = 1*, …, M*. It is supported by computer experiments as follows. Firstly, we generated random BNs and identified all basins by applying depth-first-search. Next, for a randomly selected initial state **v**^0^, we calculated the Hamming distance between **v**^0^ and all other states. Interestingly, the distribution of *d*(**v**^0^, **v**) for **v** ∈ *A*_*j*_ with a fixed *j* is found to be binomially distributed with 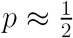 (Fig.4). Therefore, we hypothesize that 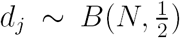. This empirical finding also supports the assumption of random partition in Theorems 1, 2, and 3.

**Fig. 4:**
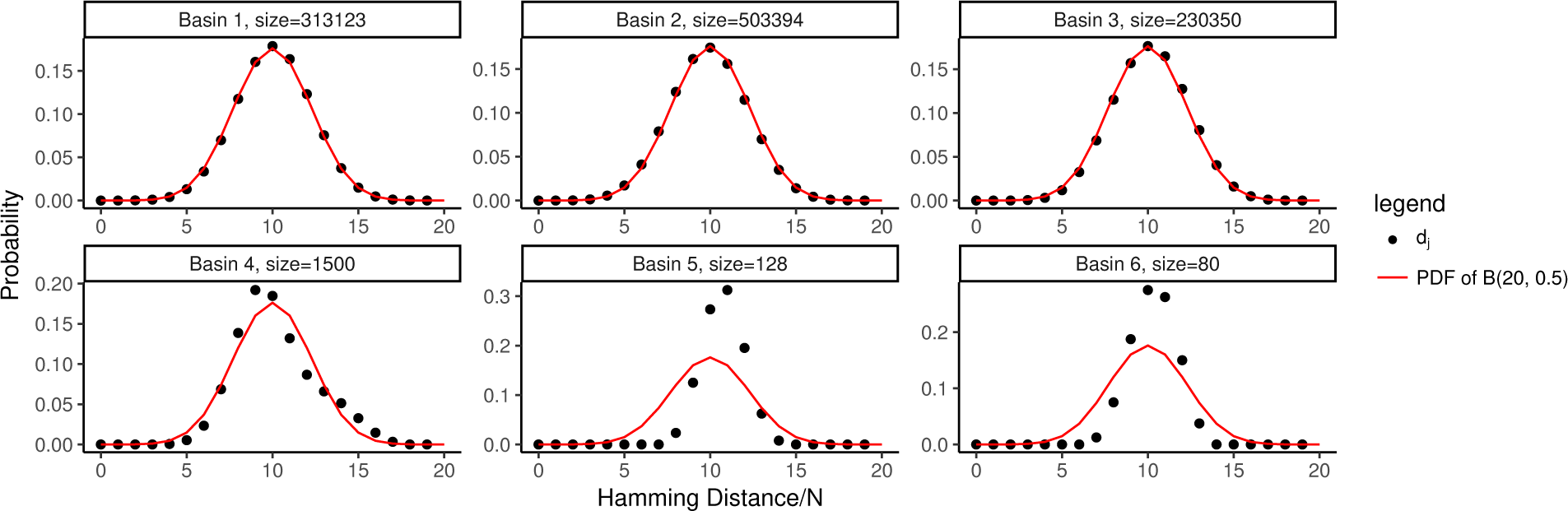
Distribution of *d*_*j*_ in a BN of *N* = 20 and *M* = 6 as an example. In each basin, the larger the basin is, the more the *d*_*j*_ follows binomial distribution with 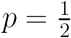.

We also found empirically that the basin size follows a power-law distribution. We examined both random BNs with *N* = 20 and *K* = 3 (maximum in-degree) and scale-free BNs of *N* = 20 and *γ* = 2.5 (in-degree distribution). The results are shown in Fig. 5, from which we can see that the basin size follows a power-law. Berdahl *et al*. had similar findings on NK-BNs [52]. Specifically, we investigated the log-log plots of the distribution of basin size and found that the slopes for those BNs with maximum in-degree (bound = 3) are −1s (error tolerance 0.05), but for those of scale-free BNs are more diverse in [−3*, −*1] with more than 60% around −2 (error tolerance 0.05). Recall that a power law distribution has a probability distribution function of ℙ[*X* = *x*] ∼ *x*^*−a*^ where *−a* is the slope of the corresponding log-log plot because log(*y*) = log(*C*) - *a* log(*x*) for a power law *y* = *Cx*^*−a*^. Therefore, we can make an estimation suggestion that the expected number of driver nodes needed to drive the BN to all basins can be estimated as *O*(log (*?*(*b*)*M* ^*b*^ ln *M*)) and *O*(log (*M* ln^2^*M*)) for BNs with the maximum in-degree *K* (*K* ≤ 3) and for those scale-free BNs, respectively.

**Fig. 5:**
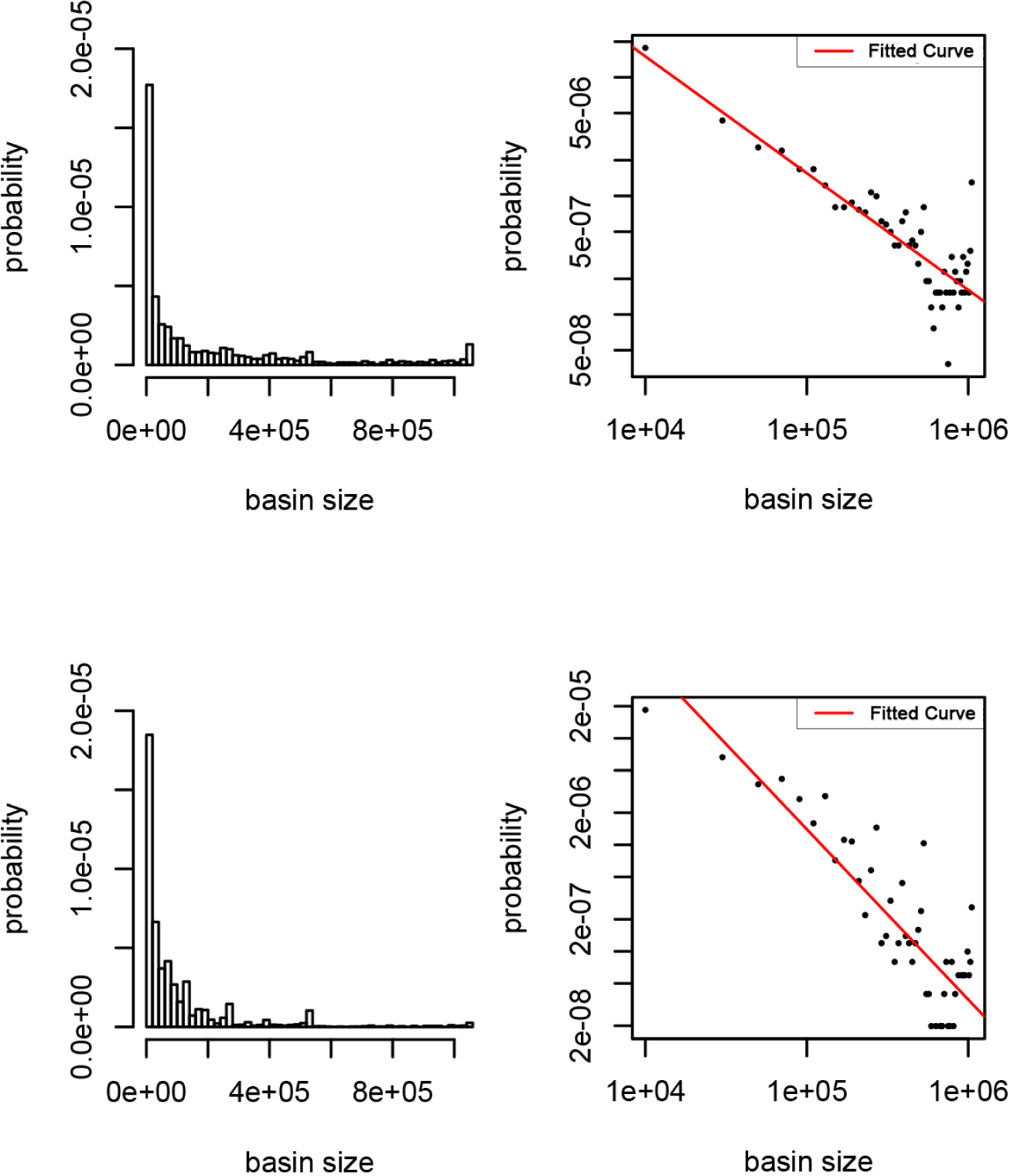
Distribution of basin size (BNs of *N* = 20 as examples). First row: on BNs with *N* = 20 and *K* = 3 (max in-degree). Second row: on scale-free BNs with *N* = 20 and *γ* = 2.5. Left: Linear scale histogram of the distribution of basin size. Right: log-log scale plot of the distribution of basin size with red lines denoting fitting lines (slope: -0.98(1st row), - 1.60(2nd row)). Data are obtained from 250 random BNs of each type.

### c. Results on the Number of Driver Nodes

Fig. 6 visualizes our key results that Attractor-dependent Control, Attractor-dependent Control, and Anycast Control are in the order of increasing the minimum control cost (i.e., the minimum number of driver nodes). Additionally, regarding the Attractor-dependent Control, we generated random BNs for 2,000 times and compared the *ξ*_(*M*)_ of each simulated BN with the expectation given by Eq. (3). We also analyzed 4 realistic BNs and calculated the corresponding value of *ξ*_(*M*)_. Results show that Eq. (3) gives reliable results of the minimum number of driver nodes for both random BNs and realistic BNs (see Fig. 6), illustrating the feasibility of taking the minimum set of driver nodes for the control of gene regulatory networks. Although the results of simulated BNs are a bit larger than the expectation given in Eq. (3) due to the constraints on in-degree bound, the error is small and tolerable. Meanwhile, the upper bounds given in Theorems 1, 2, 3 and the lower bound of log_*N*_ (*M*) serve as good boundaries for Eq. (3).

**Fig. 6:**
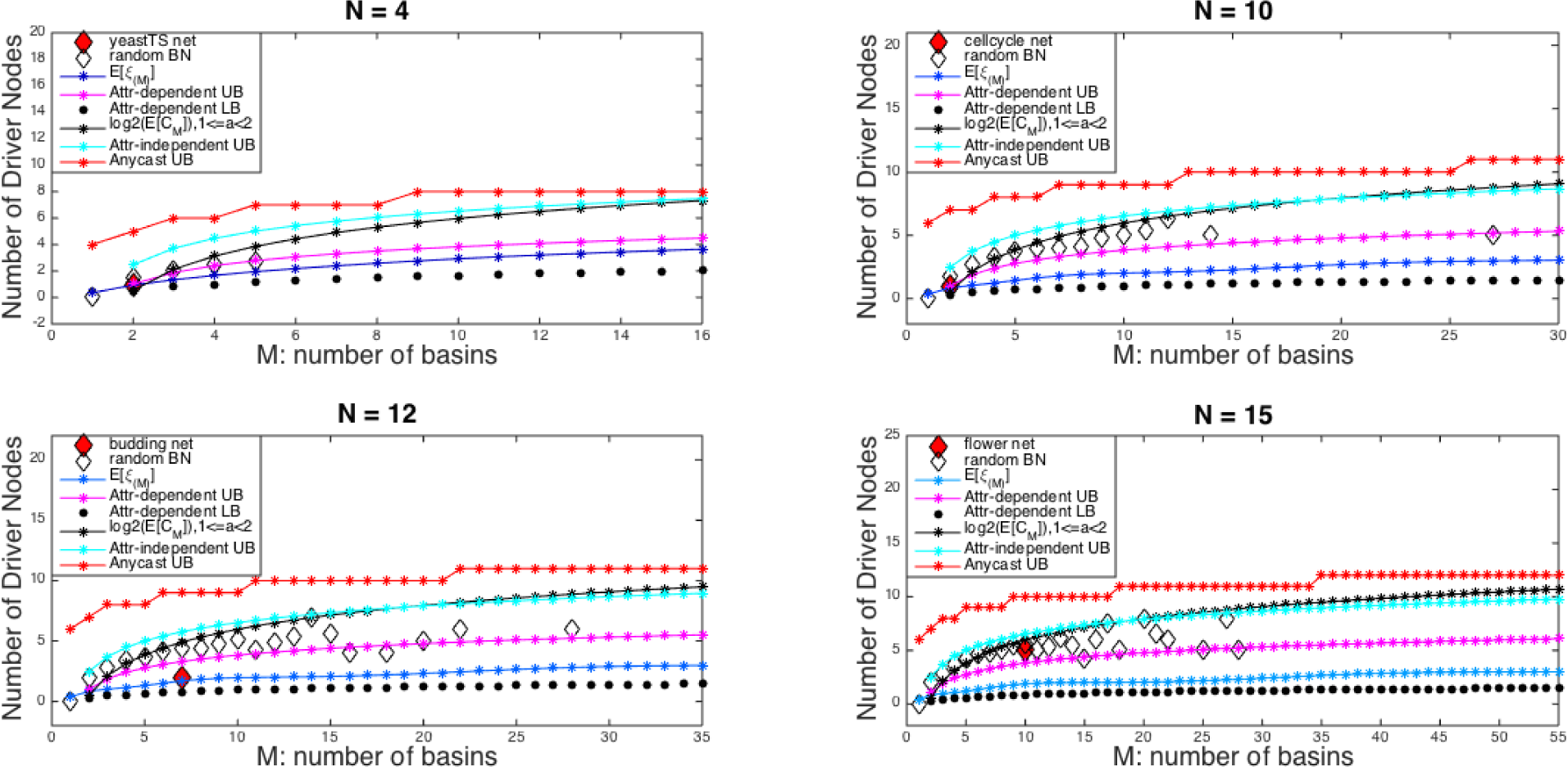
Comparison of the number of driver nodes between simulation and theoretical results. “attr”, “UB”, “LB” are short forms of “attractor”, “upper bound”, “lower bound”, respectively.

Meanwhile, it shows that CCP is a very good model to formulate this problem. Firstly, we generated random BNs. Next, we drove the BN from a random initial state to all basins with the least number of bits (denoted as *d*_*i*_ in the *i*th simulation) for 1000 simulations. The value of *d*_*i*_ was obtained as follows. For *d*_*i*_ = 1*, …, N*, we checked all combinations of *d*_*i*_ bits in **v**^0^ to see if we could drive the BN from **v**^0^ to all basins with these *d*_*i*_ bits. We returned *d*_*i*_ if it succeeded, otherwise we increased *d*_*i*_ by 1. Then we compared the average, 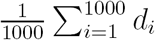, with log (𝔼[*C*_*M*_]). As shown in Fig. 7, log_2_ (𝔼[*C*_*M*_]) provides a very good prediction on the number of driver nodes (error ≤ 1), indicating that CCP is a very good model to formulate this problem. Note that although *N* is small, we are considering all 2^*N*^ states in each simulation, which we think is enough to elucidate the comparison.

**Fig. 7:**
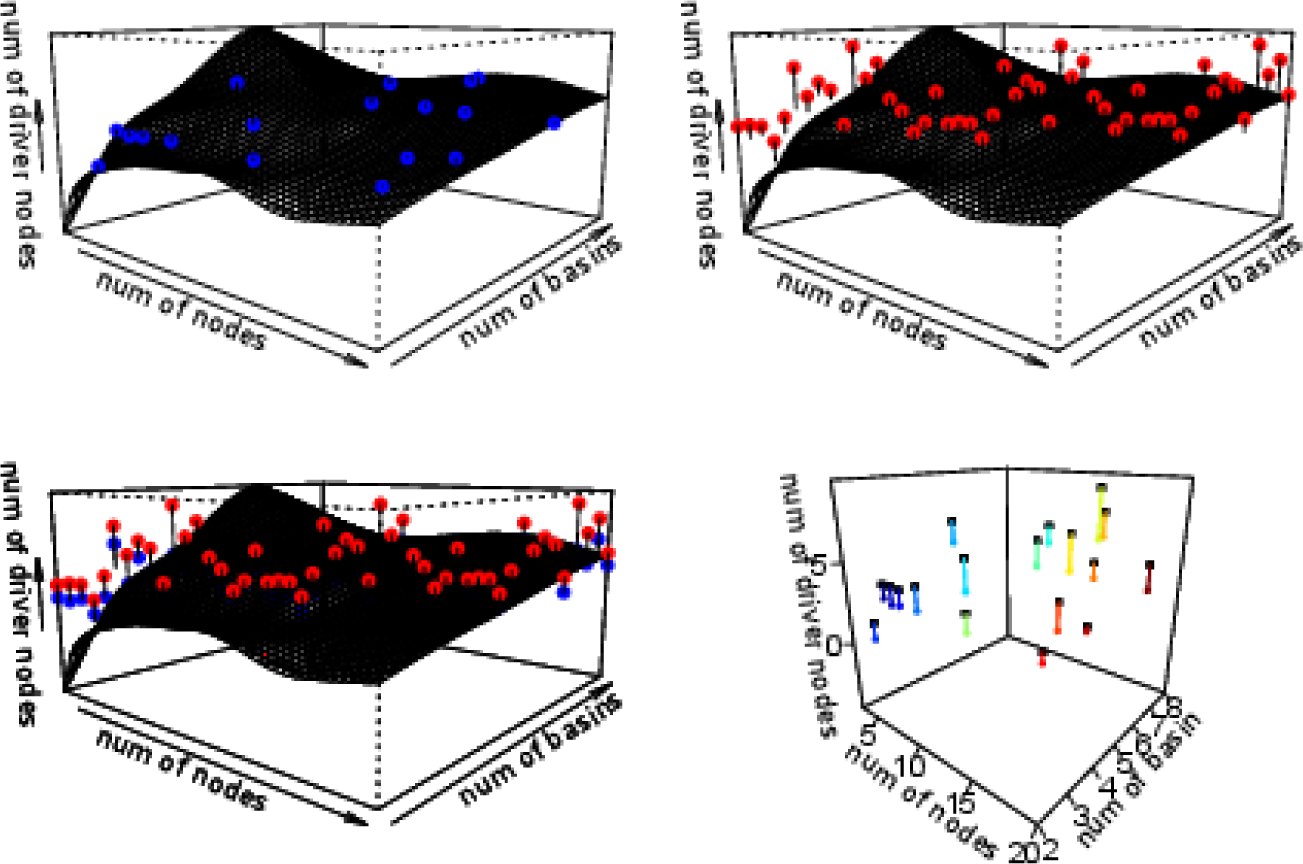
Number of driver nodes obtained by random BNs and that by CCP. First row: dot plots and fitting surfaces of simulation results (left), Eq. (4) (right). Bottom left: overlap of figures in the 1st row. Bottom right: connected by colorful segments, black squares and colorful dots denote the results obtained from simulated BNs and Eq. (4), respectively.

Recall that we give the formula of 𝔼[*ξ*_(*M*)_]. Fig. 6 shows that log_2_ (𝔼[*C*_*M*_]) provides a good upper bound of 𝔼[*ξ*_(*M*)_]. The number of driver nodes calculated by log_2_ (𝔼[*C*_*M*_]) is greater than that by 𝔼[*ξ*_(*M*)_] because the bits selected in the Attractor-independent Control have to be specified (fixed) to drive the BN to all basins but it is not in the Attractor-dependent Control.

## V. Conclusion

In this paper, we theoretically showed under a reasonable assumption that only a small number of driver nodes are required for controlling Boolean networks if the targets are restricted to attractors. This result explains previous empirical findings [38, 41] and suggests that control of biological networks might not be so difficult if the targets are steady states. We also performed computational experiments using artificial networks and realistic biological networks for verifying our theoretical findings. In addition, we pioneered the idea of formulating the minimum control cost-related problem to Coupon Collector’s Problem, which might be useful for further studies on the minimum driver set problem for other mathematical models of biological networks. The Max-min Analysis Method is utilized to get a more accurate bound in Attractor-independent Control problem, while the difficulties of applying such method in another two problems lie in the complexity of their asymptotic forms.

We focused on the cases in which control is applied only at *t* = 1. This assumption is reasonable since giving heavy controls (e.g., change of gene expression values) at many time steps is not feasible. However, if we allow control operations at multiple time steps, the obtained bounds might be improved. Such an improvement is left as an open problem. We have assumed that the structure of a BN has already been embedded in the generation of the basins of attractors, However, clarifying the relationship between the structure of a BN and the distribution of basins is also important but difficult, and thus is left as future work.

